# Can we conjointly record direct interactions between neurons *in vivo* in anatomically-connected brain areas? Probabilistic analyses and further implications

**DOI:** 10.1101/2020.12.07.415125

**Authors:** Sean K. Martin, John P. Aggleton, Shane M. O’Mara

## Abstract

Large-scale simultaneous *in vivo* recordings of neurons in multiple brain regions raises the question of the probability of recording direct interactions of neurons within, and between, multiple brain regions. In turn, identifying inter-regional communication rules between neurons during behavioural tasks might be possible, assuming conjoint activity between neurons in connected brain regions can be detected. Using the hypergeometric distribution, and employing anatomically-tractable connection mapping between regions, we derive a method to calculate the probability distribution of ‘recordable’ connections between groups of neurons. This mathematically-derived distribution is validated by Monte Carlo simulations of directed graphs representing the underlying anatomical connectivity structure. We apply this method to simulated graphs with multiple neurons, based on counts in rat brain regions, and to connection matrices from the Blue Brain model of the mouse neocortex connectome. Overall, we find low probabilities of simultaneously-recording directly interacting neurons *in vivo* in anatomically-connected regions with standard (tetrode-based) approaches. We suggest alternative approaches, including new recording technologies and summing neuronal activity over larger scales, offer promise for testing hypothesised interregional communication and source transformation rules.

## 1 Introduction

The technique of choice to understand neuronal circuit dynamics *in vivo* continues to be intracranial recordings of single and multiple neurons (and of associated signals, such as local-field potentials). Recent technical advances in neuronal recording methodologies (such as Neuropixels probes; Jun et al. 2017; Steinmetz et al. 2020) allow recording of large numbers of single neurons within and between multiple brain regions simultaneously (Siegle et al. 2019; Stringer et al. 2019; Steinmetz et al. 2019; Allen et al. 2019). These approaches offer a level of analysis previously impossible using tetrode-based recordings. Stringer et al. (2019), for example, simultaneously-recorded *∼*3,000 neurons across the brain using Neuropixels and *∼*10,000 neurons in visual cortex using calcium imaging, of which an unknown proportion might be directly interconnected. However, these recordings remain spatially- and temporally-incomplete samplings, and achieving a true population recording is an unrealistic goal (Buzsáki 2004). These methods undersample the activity of the true number of neurons present within brain regions, as the number of neurons present (and potentially active) is orders of magnitude greater than the number of recording sites available.

Large-scale, inter-regional, conjoint recordings leave the problem of understanding granular information processing rules between neurons in connected brain regions. Many previous studies of inter-areal interactions have analysed spiking activity of pairs of neurons in different areas (Semedo et al. 2019). The probability of connection between pairs of neurons at a given distance from each other, usually on the scale of a few hundred micrometres, has previously been considered (e.g. Liley and Wright 1994; Braitenberg and Schüz 1998; Holmgren et al. 2003; Kalisman, Silberberg, and Markram 2003). When considering inter-areal interactions at the neuronal level, it is important to consider the proportions of sampled neurons that might interact with each other in each recording, and how direct this interaction is. Here, we provide a method for estimating proportions of sampled neurons in anatomically-connected brain regions.

Structural connectivity provides the substrate for information transfer in the brain (Goulas, Uylings, and Hilgetag 2017). Understanding source transformation rules may enable us to probe how downstream neuronal observers take advantage of this information (Buzsáki and Tingley 2018). In other words, what are the source transformation rules applied to given inputs from area A to area B, and can we reasonably expect to infer them, given the sampling that current methodologies afford? Being able to answer this question will allow us, in turn, to approach the question of how transformations of neural coding within and between brain regions might eventuate in the generation of behaviour, and provide a mechanistic basis for understanding cognition.

We consider the following question, using an inductive, analytic, approach: given imperfect samplings, how likely are we to record direct interactions between neurons in different brain areas? We focus on connections between neurons, and not synaptic weights, firing patterns and so on, because structural connectivity provides the basis for any communication between connected brain regions. These latter parameters likely provide the source transformation rules between connected brain regions, but depend, in principle, on the pattern of connectivity between the regions.

We consider several neurons in multiple brain regions, with simplified connections between neurons in these regions (see Figure 1). We use the hypergeometric distribution, which describes the probability of sampling items with a particular property from a population without replacement; in this case, recording from neurons sending or receiving connections from a population of neurons. From the hypergeometric distribution, we derive the probability distribution for the possible number of interacting neurons that might be recorded between the populations of neurons. This mathematically-derived distribution and estimation of the number of connected neurons is validated by Monte Carlo simulations of directed graphs. We apply this to illustrative examples using rat hippocampal and subicular cell counts, and examples from the Blue Brain Project’s model of the mouse neocortex (Reimann et al. 2019). Our results suggest the probability of conjointly recording directly-interacting neurons may approach zero for tetrode recordings, but recordings of directly-interacting neurons may be feasible with current high-site count recording technologies in well-connected brain regions. This latter possibility emphasises the need for new methods to detect and analyse such conjoint recordings.

**Figure 1:**
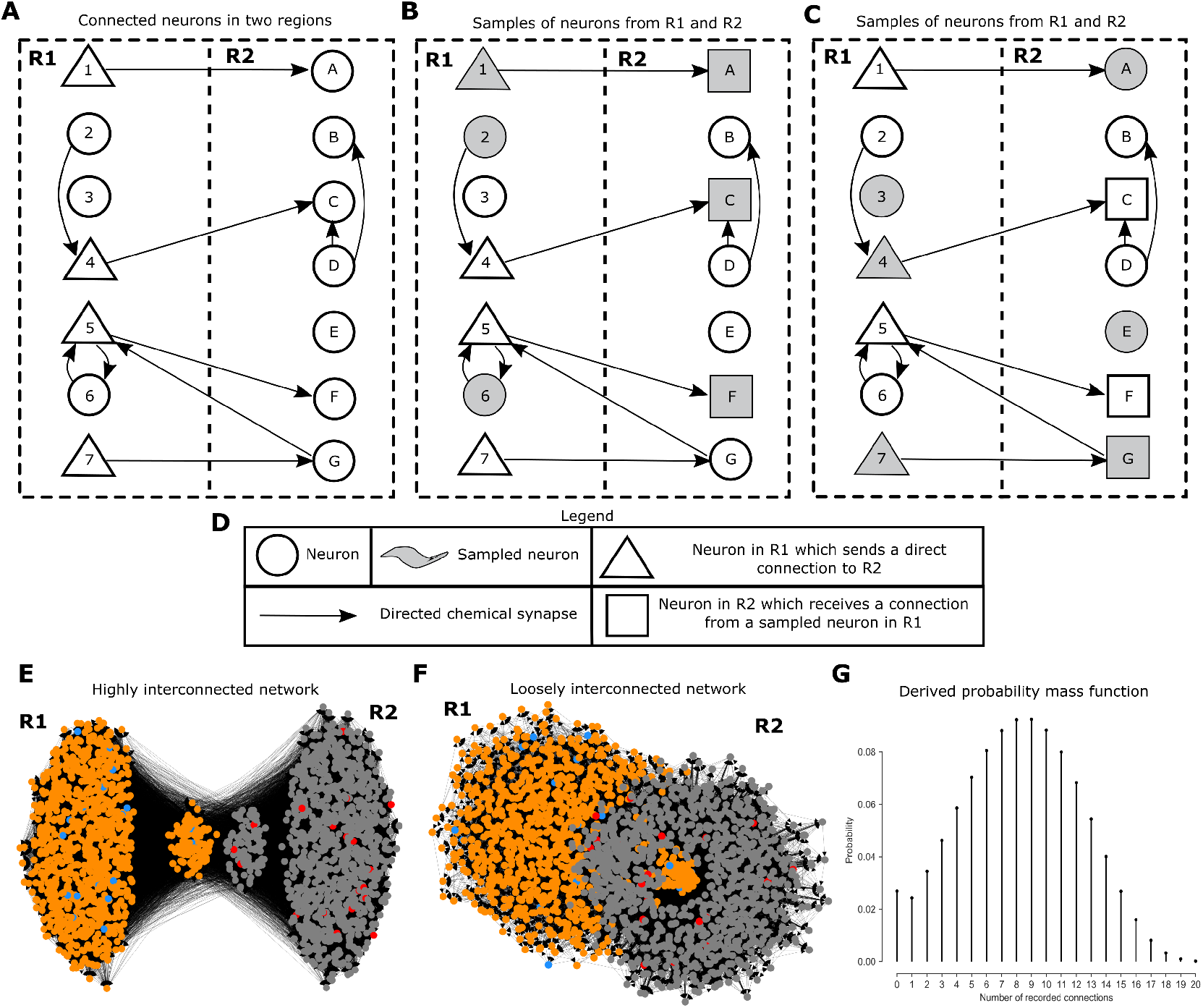
Problem illustration recording connections between sampled neurons. (A) Connections between neurons in two regions, where the neurons in R1 which send direct connections to R2 are marked. (B) Randomly sampling three neurons from R1 and three neurons from R2 results in differing numbers of sampled neurons which are connected. In this case, all three of the sampled neurons in R2 receive a connection from a neuron sampled in R1, but two are indirect connections. (C) As in (B), but one of the sampled neurons receives a connection from a neuron sampled in R1. (D) Legend for panels A, B, and C. (E, F) Examples of larger-scale networks of neurons with one-thousand neurons in each region. Blue and red dots indicate representative samples of neurons in R1 and R2. (G) From the network connectivity, we derive the probability distribution of the number of sampled neurons in R2 which receive a direct connection from a neuron sampled in R1 when randomly sampling 20 neurons from R1 and R2 in (F).

## 2 Results

Given a model *M* for the direct synaptic connections within and between two brain regions *A* and *B*, this method estimates the probability distribution of the number of neurons recorded in *B* that receive a connection from a neuron recorded in *A*, when randomly sampling *k* neurons from *A* and *m* neurons from *B*.

### Box 1

**Model assumptions**

1. Autapses are not considered.
2. Multiple connections between two specific neurons are considered as one connection. The direct influence of one neuron onto another is typically weak, being mostly represented by a single synapse (Braitenberg and Schüz 1998).
3. Synaptic connection strength is not considered - just the presence or absence of a synapse.
4. The problem is considered as a fixed state - or point estimate - so, for instance, the dynamic nature of brain connectivity over time during chronic recordings is not considered.
5. We do not distinguish between inhibitory and excitatory connections. Setting up the model with statistics of, say, only inhibitory connections would then give probabilities for that type of connection.
6. Only chemical synapses are considered and not electrical synapses (gap junctions), because electric synapses could be considered as undirected connections, while chemical synapses are directed (Reimann et al. 2017).
7. Neurons are uniformly sampled from the areas in question, and topographical changes or gradients in the area are not considered. For instance, during simultaneous recordings in rat CA1 and subiculum, the model should be set up with appropriate statistics relative to the recording device placement and angle of approach, as, for example, the projections from proximal CA1 to distal subiculum and distal CA1 to proximal subiculum are different.
8. We do not consider the issue of the probability that spiking from a neuron in area *A* causes spiking in a neuron in area *B* (they are likely to be very weakly interacting); we just consider that they are anatomically-connected.

### 2.1 Description as a directed graph

Consider recording *k* and *m* neurons respectively from two brain regions or areas *A* and *B*, each containing *N* and *M* neurons overall. The regions *A, B* are two disjoint sets of vertices, representing neurons, in a directed graph *G* = (*V, E*), describing the physical connectivity of the neurons through directed chemical synapses. *E*, the set of arrows connecting the vertices, indicates a direct connection between two neurons in *V* (Reimann et al. 2017), such that *v*_*i*_, *v*_*j*_ *∈ E* indicates a synaptic connection between *v*_*i*_ and *v*_*j*_ where *v*_*i*_ is the presynaptic neuron, and *v*_*j*_ is the postsynaptic neuron.

Our assumptions impose the following constraints on the graph. The graph is simple directed (6 - chemical synapses); it has no loops (1 - a neuron is not connected to itself) and at most one arrow exists with the same source and target nodes (2 - multiple synaptic connections between two neurons are bundled into one connection). Furthermore, the graph is static (4 - nodes and edges are fixed), and all nodes and edges in the graph are of a single type (3, 5 – no synaptic weights, no inhibitory/excitatory split).

Our primary question can be restated as: *what is the probability that a random sample of vertices V*_*S*_ *∪ V*_*E*_ *from disjoint regions (V*_*S*_ *⊂ A, V*_*E*_ *⊂ B, A ∩ B* = ∅*) contains directed paths in which x vertices in V*_*E*_ *are reachable from V*_*S*_.

### 2.2 Calculating connection probability

Let *X* be the random variable whose value is the number of neurons recorded in *B* which receive at least one connection from a neuron recorded in *A*. Furthermore, let *X*_*A*_ be the random variable whose value is the number of neurons recorded in *A* sending a connection to *B*, and *X*_*B*_ be the random variable whose value is the number of neurons in *B* receiving a connection from a neuron recorded in *A*. By the chain rule of probability, and taking the marginal distribution over *X*_*A*_ and *X*_*B*_, the probability of recording *x* such neurons in *B* is then

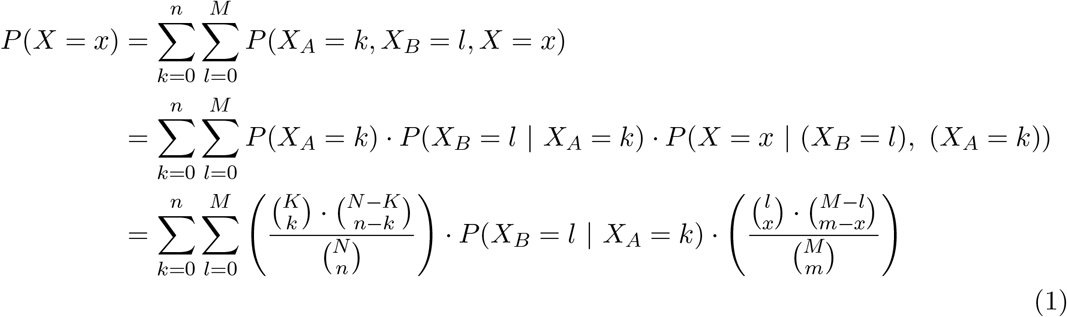

where:

*N* = number of neurons in *A*

*n* = number of neurons recorded in *A*

*K* = number of neurons in *A* which send connections to *B*

*k* = number of recorded neurons in *A* which send connections to *B*

*M* = number of neurons in *B*

*m* = number of neurons recorded in *B*

*l* = number of neurons in *B* which receive connections from neurons recorded in *A*

In the above formula, two things are of note. Firstly

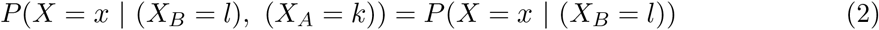

since knowing how many sampled neurons in *A* send connections to *B* is redundant to calculating *P* (*X* = *x*) if the number of neurons in *B* which receive connections from the neurons sampled in *A* is known. Secondly, *P* (*X*_*A*_ = *k*) and *P* (*X* = *x* | (*X*_*B*_ = *l*)) are both calculated from the hypergeometric distribution, which describes the probability of sampling items with a particular property from a population without replacement; in this case, recording neurons sending or receiving connections from a population of neurons.

### 2.3 Estimating the proportion of receivers

Calculating the above distribution requires calculating *P* (*X*_*B*_ = *l* | *X*_*A*_ = *k*), the probability of *l* neurons in *B* receiving at least one connection from a neuron recorded in *A*, given that *k* neurons are recorded in *A* which send direct connections to *B*. Due to the indeterminacy of which neurons would be recorded, the possibility of recording neurons in *A* which project to thousands of neurons, or just a couple of neurons in *B* must be considered. This calculation is where the connectivity information between the two regions is required. For example, the Blue Brain Project’s recipe for how connections are made between regions (Reimann et al. 2019).

Firstly, we limit the maximum geodesic distance considered between nodes in *A* and *B* in the graph, or the number of synapses that can be used to connect the neurons in *A* and *B*, to a value *D*. Building up the interactions of outgoing (*AB*), recurrent (*BA*), and local (*AA, BB*) connections grants the full picture, as these building blocks are used to form any paths between *A* and *B*. Given *D*, the sum over 2^*D*^ *−* 2 variables must be taken to compute the marginal distribution *P* (*X*_*B*_ = *l* | *X*_*A*_ = *k*).

Let *X*_*AB*_ be the random variable denoting the number of neurons recorded in *B* directly connected to a neuron recorded in *A*. Similarly, let *X*_*AAB*_ denote the number of neurons *B* not receiving a direct connection from a neuron recorded in *A*, but receiving a connection from a neuron recorded in *A* via two synapses, where the first synapse remains in *A*, and so on. Then, for the max geodesic distance *D* = 1 (direct connections only)

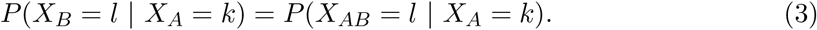

While for the max geodesic distance *D* = 2 (two synapses allowed), the sum is taken over two variables

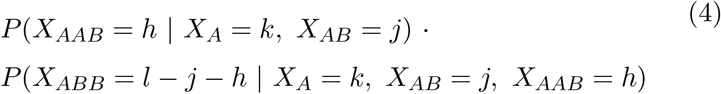

Consider computing *P* (*X*_*B*_ = *l* | *X*_*A*_ = *k*) for direct connections. We assume the random variables *R*_1_, *R*_2_, … , *R*_*k*_, representing the number of synapses that each sampled neuron in *A* that projects to *B* sends to *B*, (only neurons in *A* which send synapses to *B* are considered, as otherwise, the random variable takes the value 0 with probability 1) are independent and identically distributed with mean *µ* and variance *σ*^2^. Then *S*_*k*_ = *R*_1_ + *R*_2_ + … + *R*_*k*_ represents the number of outgoing synaptic connections from *A* to *B*. By the central limit theorem *S*_*k*_ = *R*_1_ + *R*_2_ + … + *R*_*k*_ *≈ N* (*kµ, kσ*^2^) for high values of *k*. For *k* below these values, a recursive convolution operation is applied to the distributions of the random variables (Grinstead and Snell 2012, see Methods). Finally, we apply a function *f* to *S*_*k*_ to produce a random variable *Y* = *f* (*S*_*k*_) whereby *P* (*Y* = *l*) = *P* (*X*_*B*_ = *l* | *X*_*A*_ = *k*), the original distribution we aimed to compute. The function *f* (*s*) computes the expected number of unique neurons connected to in *B* with *s* synapses from *A* to *B*, and is defined as follows; given *M* neurons in *B*

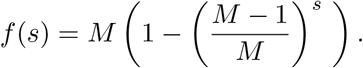

### 2.4 Mean estimation

Calculating *P* (*X* = *x*) using the full distributions of *P* (*X*_*B*_ = *l* | *X*_*A*_ = *k*) for each *k* can be too time consuming, so instead, we can consider the random variable *X*_*B*_ to be distributed as a single point at the mean of the true distribution with 0 variance, whereby *P* (*X*_*B*_ = *l* | *X*_*A*_ = *k*) = 1 if *l* = *µ*, and 0 otherwise. This reduces the accuracy, but increases the speed, of calculating *P* (*X* = *x*). This approach is generally necessary for large graphs (*>* 100,000 nodes) with a maximum geodesic distance greater than 1 (see Supplemental material).

### 2.5 Topography of brain regions

The probability of recording anatomical connections between neurons depends on the physical properties of the neural recording device (e.g. tetrodes sample in spheres; probes in capsules; calcium imaging in thin layers) and angle of approach (e.g. parallel to a columnar structure or cutting across columns). Here, we do not directly model space, instead assuming a uniform sampling in the areas being considered. With good knowledge of the region’s topography, and confidence that the placement of the recording device is approximately correct, a ‘constrained’ version of the statistical model can be established, whereby the statistics reflect relative positioning of the recording probe and underlying anatomical connectivity. However, in cases where underlying topographies or gradients are very loose or difficult to discern (e.g. CA1 and subiculum projections to prelimbic cortex), or the topographies are too cumbersome to model, the recording probe can be assumed to uniformly sample from the entire brain area in question. These considerations stress the importance of anatomical sophistication and guidance in deciding placements of in vivo recording devices.

#### Box 2

**Example calculation of a connection distribution**

Consider the situation described Figure 1. Here, there are 7 neurons in each region; there are 4 neurons which send exactly one direct connection from the first region, and we randomly sample 3 neurons in each region. Let *X* denote the random variable whose value is the number of sampled neurons in the second region receiving a direct connection from a neuron sampled in the first region. A full calculation of *P* (*X* = *x*) is below for this case:\

Let

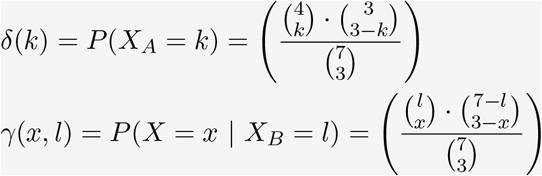

Then

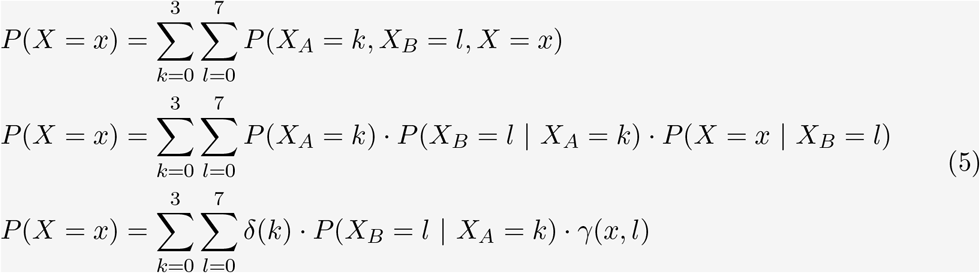

With

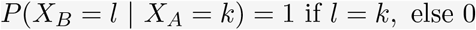

Finally

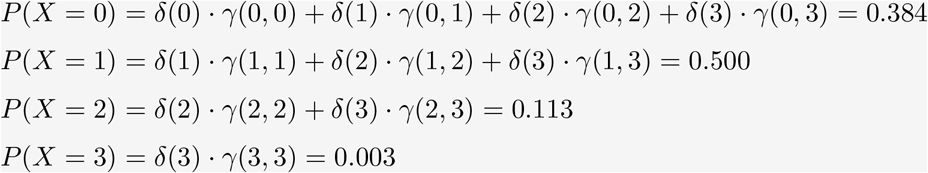

### 2.6 Case study: Rat hippocampal formation

Here, we draw on previously published estimates of the numbers of neurons in differing regions of the rat hippocampal formation, with a specific focus on areas CA3, CA1 and subiculum, as these, respectively, receive large, well-defined, anatomical projections from the preceding area.

We present results for CA3 to CA1 (see Figure 2), where key statistics are available from neuroanatomical studies (see Table 1). Consider two situations here; recording from a tetrode in CA1 and CA3, as well as a neuropixels probe. A tetrode is assumed to sample from a uniform sphere, while a neuropixels probe is assumed to sample from across the whole brain region for simplicity. In both cases, it is assumed that the topographical organisation of neurons within these spaces is uniform.

**Table 1:** Rat hippocampal formation neuron estimates and assumptions

| Source | Result |
| --- | --- |
| Andersen et al. 2006 | 390,000 and 290,000 principal neurons in CA1 and subiculum, respectively. |
| Amaral, Ishizuka, and Claiborne 1990 | CA3 contains 303,930 pyramidal cells. |
| Bezaire and Soltesz 2013 | 89% (347,100) of neurons in rat CA1 are pyramidal cells. |
| Ropireddy, Bachus, and Ascoli 2012 | CA1 occupies a volume of $18.8\text{mm}^3$, CA3 $12.2\text{mm}^3$. |
| West, Danscher, and Gydesen 1978 | The subiculum occupies a volume of $10\text{mm}^3$. |
| Amaral, Ishizuka, and Claiborne 1990 | A CA1 pyramidal cell might receive input from about 1.8% of the CA3 pyramidal cell population. |
| Arszovszki, Borhegyi, and Klausberger 2014 | Pyramidal cells in ventral CA1 commonly have terminals in ventral subiculum, and some projections bifurcate. |
| Reimann et al. 2019 | There is a distribution of synapse projections around the mean. |
| Mechler et al. 2011 | A tetrode samples neurons in a $130\mu\text{m}$ radius sphere (volume $0.0092\text{mm}^3$). |
| Gray et al. 1995 | A tetrode in visual cortex has a mean yield of about 5 cells. |

**Figure 2:**
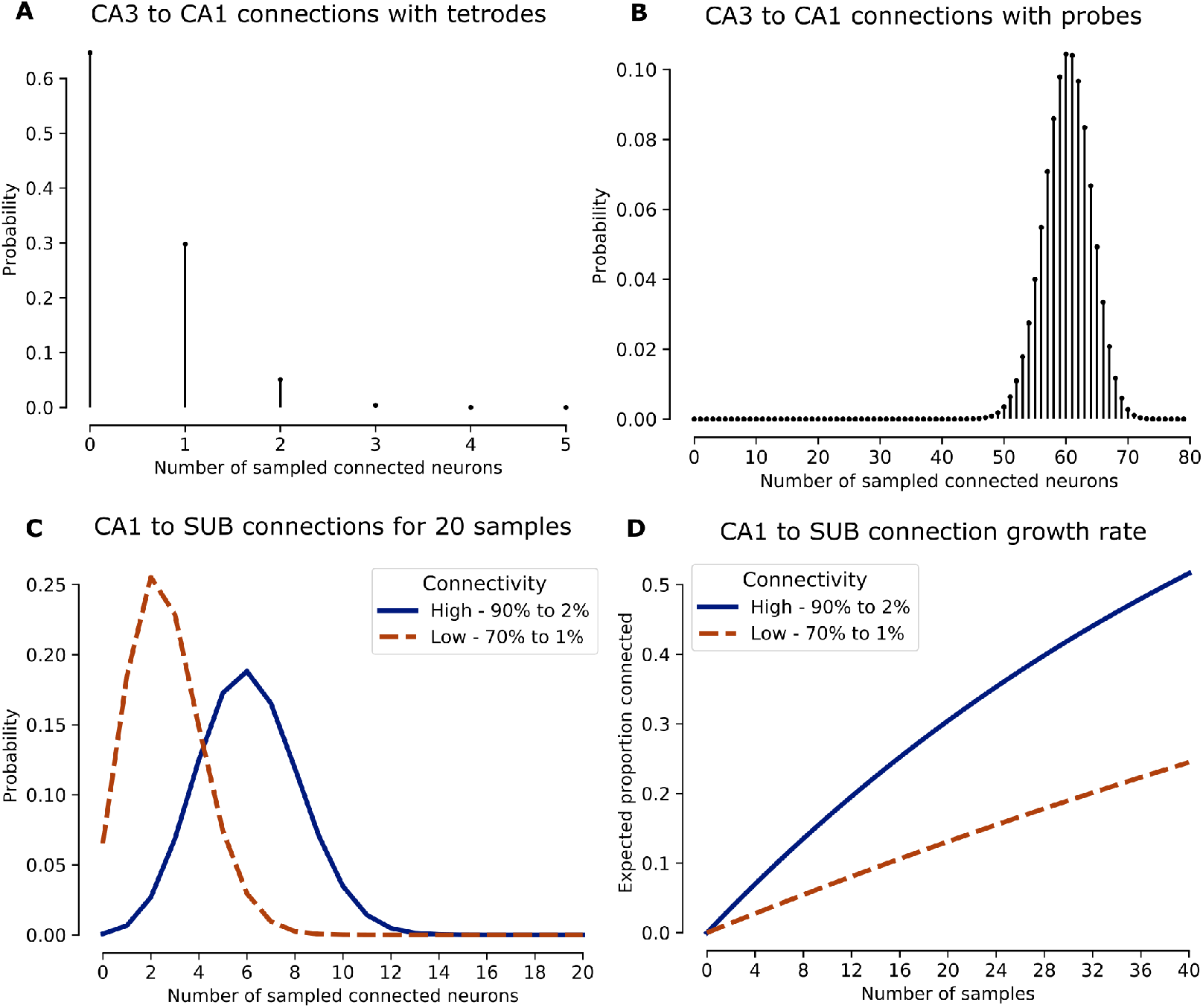
Modelling the probability of simultaneously recording directly connected neurons in rat CA3, CA1, and the subiculum. (A) With a tetrode in both rat CA3 and CA1, there is only a 36% chance of simultaneously recording any directly connected neurons. (B) Recording directly connected neurons in rat CA3 and CA1 with 79 samples (the average of a neuropixels probe Jun et al. 2017) is more likely, with a 95% chance of between 67% and 86% of neurons recorded in CA3 receiving a direct connection from at least one neuron in recorded in CA1. (C) With 20 samples simultaneously obtained from rat proximal CA1 and distal subiculum, the connected neurons are highly dependent on the underlying connectivity. If 90% of proximal CA1 pyramidal cells project to 2% of the distal subicular pyramidal cell population, *∼*6 connected neurons are expected, but if instead, 70% project to 1%, then *∼*2.5 connected neurons are expected. (D) Similarly to (C), the growth rate of connections is highly dependent on the underlying connectivity between rat proximal CA1 and distal subiculum.

With a tetrode (5 neurons) in both rat CA3 and CA1, there is just a 36% chance of simultaneously recording any directly connected neurons, while for a probe (79 neurons) there is 95% chance of between 67% and 86% of neurons recorded in CA3 receiving a direct connection from at least one neuron in recorded in CA1 (see Figure 2). This is calculated from (see Supplemental material for further details):

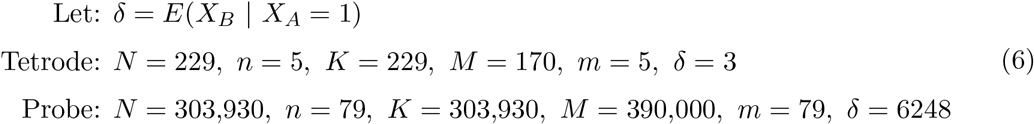

Where *δ* = *E*(*X*_*B*_ | *X*_*A*_ = 1) denotes the expected number of neurons in CA3 that receive a direct connection from a single neuron in CA1. For instance, with the probe, we then assume that a single CA3 pyramidal cell sends an average of 6305 random synapses to CA1 pyramidal cells, so a single CA3 pyramidal cell would be expected to connect to 6248 CA1 pyramidal cells (more than one synapse can project to the same CA1 neuron).

The subiculum is the primary output structure of CA1 (Amaral and Witter 1989; Witter and Groenewegen 1990; O’Mara et al. 2001); CA1 pyramidal cells mostly differ along the dorsal-ventral axis and are more homogenous along the proximal-distal axis (Ishizuka, Cowan, and Amaral 1995; Cembrowski et al. 2016). CA1 pyramidal cells project in a columnar fashion to subiculum (Amaral, Dolorfo, and Alvarez-Royo 1991), which is segregated into discrete subclasses (Cembrowski et al. 2018). CA1 neurons display a relatively homogenous place code, whereas subicular neurons have a very heterogenous spatial code (Brotons-Mas et al. 2017). Consider recording pyramidal cells in the rat proximal CA1 (near CA3), which project to distal subiculum (Amaral, Dolorfo, and Alvarez-Royo 1991).

Problematically, for subicular projections from CA1, we are uncertain about many of the key numbers (distribution of projections, number of senders, proportion of pyramidal cells), as the relevant data do not exist in the literature. We, therefore, provide two scenarios, based on CA1 as follows: (for simplicity, we assume that one third of subicular cells lie in distal subiculum (96,666 cells), and similarly for proximal CA1 (115,700 pyramidal cells)).

- There is a high rate of projection; say, 90% of pyramidal cells in proximal CA1 project to distal subiculum, and those that do, project to 2% of the distal subicular pyramidal cell population (akin to CA3 to CA1 projection rates). Additionally, the percentage of subicular pyramidal cells is akin to CA1, roughly 90%.
- Alternatively; assume that the rate of projection is lower than CA3 to CA1, say 70% of pyramidal cells in proximal CA1 project to 1% of the distal subicular cell population. Additionally, assume the subiculum has a lower proportion of pyramidal cells than CA1, say 80%.

The results are highly dependent on the underlying connectivity (see Figure 2). For instance, if 20 samples are simultaneously obtained from rat proximal CA1 and distal subiculum, with the high rate of projection, then about 6 connected neurons are expected, but with the lower projection rate, this falls to *∼*2.5 connected neurons. This is calculated from (see Supplemental material for further details):

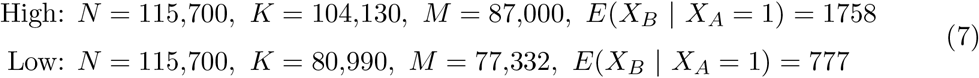

Where *E*(*X*_*B*_ | *X*_*A*_ = 1) denotes the expected number of neurons in distal subiculum that receive a direct connection from a single neuron in proximal CA1 that projects to distal subiculum.

The results suggest that in well-connected regions, conjoint recordings of directly interacting neurons could be possible, though large sample sizes are required to detect them. We estimate that roughly 40 simultaneous samples are required from rat CA3 and CA1 to expect close to 50% of the recorded population in CA3 to directly interact with at least one of the recorded CA1 neurons. Recent advances in recording technology, such as Neuropixels probes, certainly allow for this possibility. Furthermore, situations like CA1 to subiculum are conducive of simultaneously recording connected neurons as most CA1 neurons project to a relatively large subset of the subicular cell population (and the precise location of recording probe will matter too).

### 2.7 Case study: Neocortex connections

Reimann et al. (2019) present a null model of a full mouse neocortex with draft parametric rules for short- and long-range connections to stochastically connected, morphologically detailed neurons in a three-dimensional volume representing the full mouse neocortex (the authors provide full connectivity matrices between brain regions in the mouse neocortex). We apply our graph simulations and statistical estimations to this model of the mouse neocortex using these connectivity matrices (see Figure 3), with the assumption that neurons are recorded at random in these neocortical brain regions. This is both an interesting case study, and highlights the accuracy of the statistical estimation on very large graphs. For certain areas of the mouse neocortex, such as the auditory cortex, this analysis suggests that conjoint recordings of directly connected neurons are feasible with new recording technology. However, even for regions with lower connectivity, such as MOp (primary motor cortex) to SSp-ll (primary somatosensory area associated with lower limbic function), the number of sampled neurons in SSp-ll which receive connections from sampled neurons in MOp grows rapidly if the analysis method is agnostic to the number of synaptic jumps between the neurons (which is likely why techniques such as optogenetics can be so impactful, even if they act on a very small subset of the full population).

**Figure 3:**
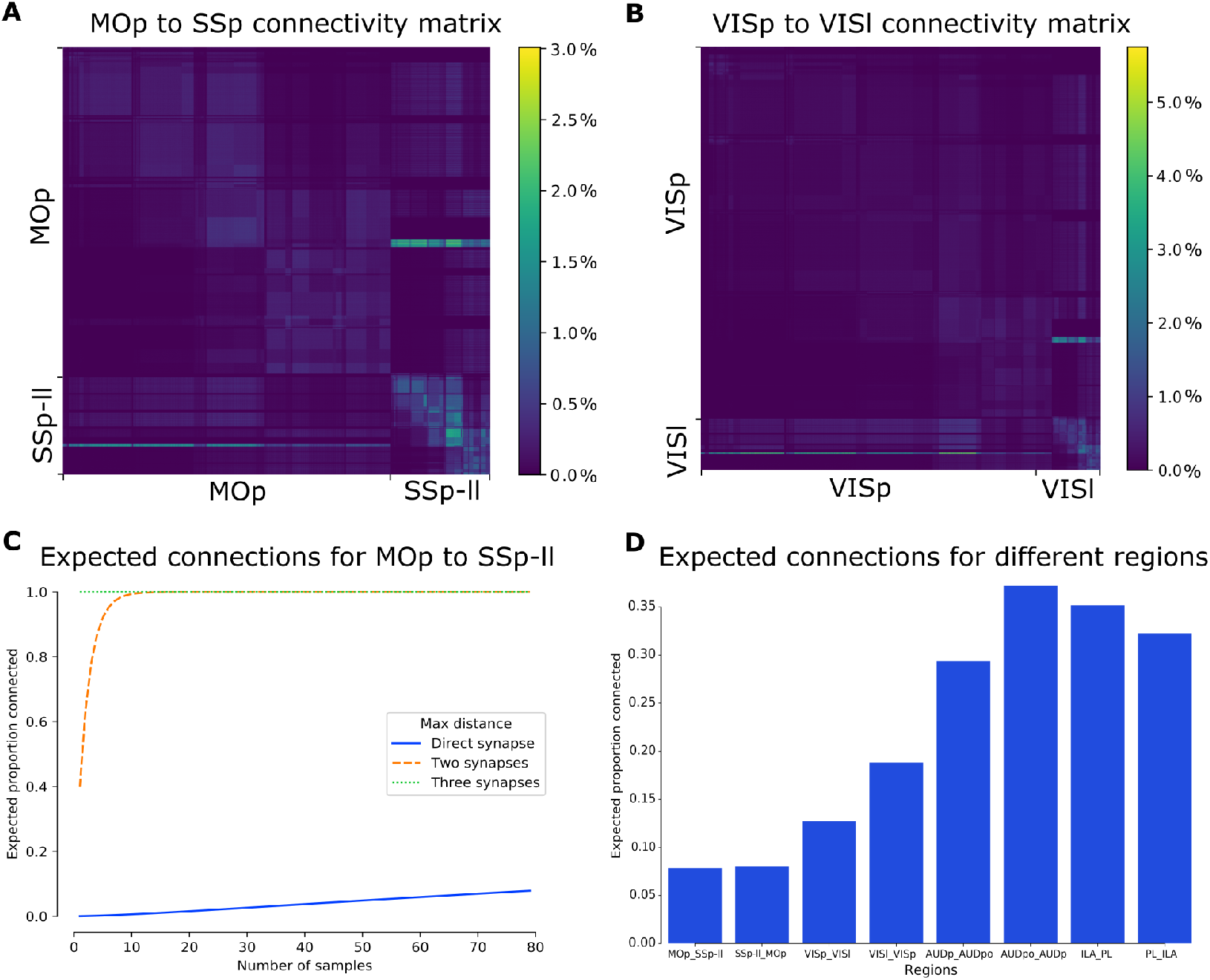
Analyses based on the Blue Brain Project’s model of mouse neocortex. (A, B) Visualising the connection matrix between brain regions in the neocortex (density of connections in 150 by 150 cubes of the full connectivity matrix is shown). The row in the matrix indicates the presynaptic site, while the column indicates the postsynaptic site. (A) Ipsilateral and local right hemisphere connections between MOp (primary motor cortex) and SSp-ll (primary somatosensory area associated with lower limbic function). (B) Ipsilateral and local right hemisphere connections between VISp (primary visual area) and VISl (lateral visual area). (C) The proportion of expected connections dependent on the number of samples recorded in each region; direct connections are close to linear, connections along at most two synapses close to exponential, and each sampled neuron in SSp-ll receives a connection from MOp along three synapses. (D) Calculating the expected proportion of direct connections for different regions with 79 recorded neurons in each region (the average of a neuropixels probe, Jun et al. 2017); *A B* indicates *A* sending to *B*. (AUDp, AUDpo - primary and posterior auditory areas; ILA, PL - infralimbic and prelimbic areas.)

### 2.8 Case study: Accuracy of statistical estimation

The accuracy of the statistical estimation is validated by a close match to simulated experiments (see Figure 4). Our statistical formula to estimate the underlying probability distributions and expectations gives similar results to those from Monte Carlo simulations of the graphs in all cases, verifying the formula’s accuracy. Of note is the close match between the simulations on the Blue Brain Project’s model of MOp to SSp-ll connectivity and our formula, indicating the statistical estimation remains accurate even when handling very large graphs with high variance. However, as the maximum geodesic distance increases, the accuracy begins to decrease (see Supplemental material). Nonetheless, having an accurate statistically-based formula to calculate expected connections and avoid costly computations is a huge benefit.

**Figure 4:**
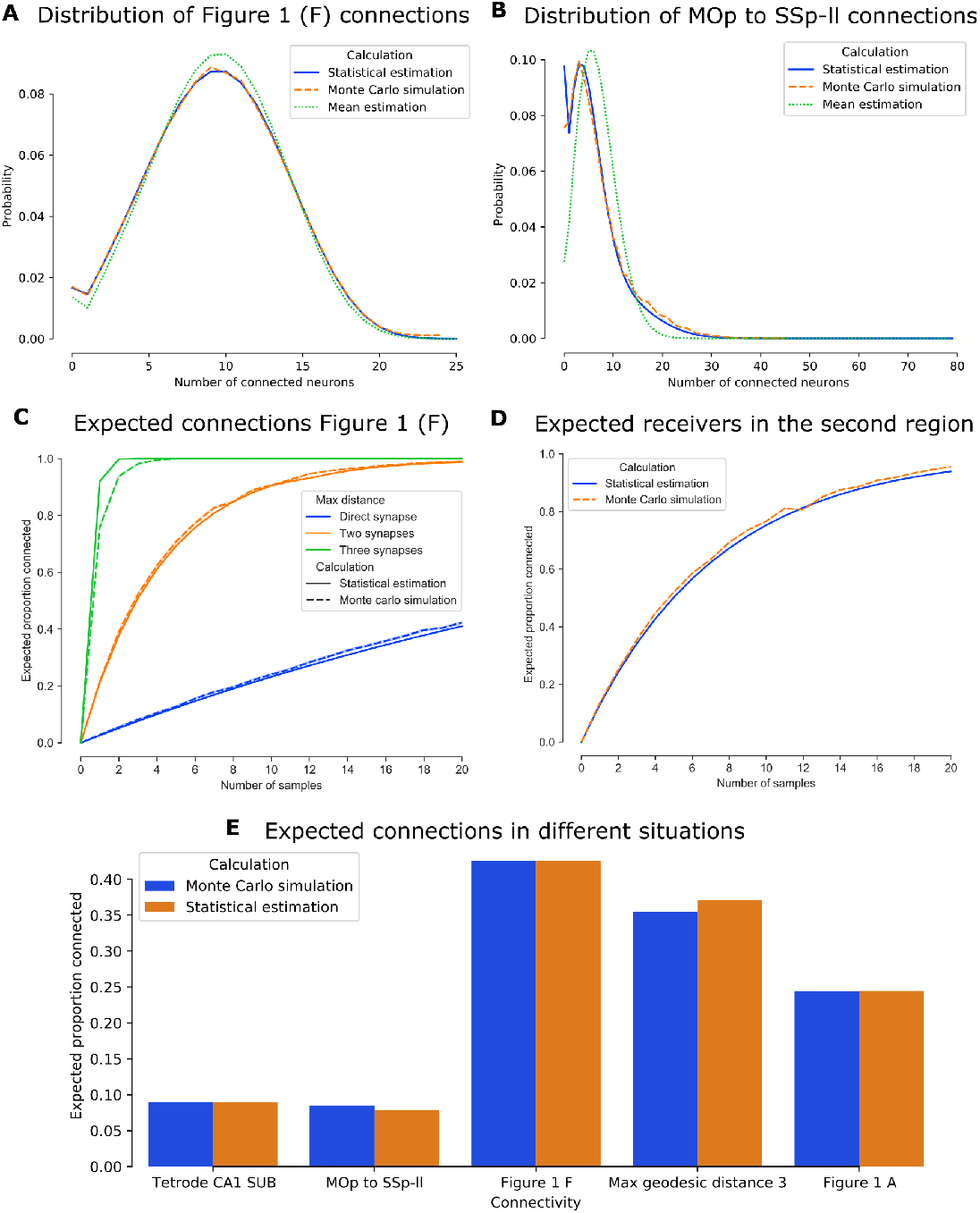
Accuracy of the statistical estimation. (A) The probability mass function is accurate, but values differ more if only using the mean to estimate (1000 simulations of 100 random graphs like the network in Figure 1, F; 20 samples in *A*, 25 in *B*). (B) As in (A), but on MOp to SSp-ll in mouse neocortex Reimann et al. 2017 with 79 samples (50,000 simulations). (C) The expected number of connections is accurate (10,000 simulations of 10 random graphs like the network in Figure 1, F). (D) A fixed number of samples *k* which send connections are taken from a network like Figure 1, F. The expected number of connected neurons in *B* is estimated from the distribution of *P* (*X*_*B*_ = *l* | *X*_*A*_ = *k*) (10,000 simulations each). (E) Assessing the expected number of connections from different types of connection strategy and brain regions (50,000 simulations each).

## 3 Discussion

A profound and unresolved issue in neuroscience is understanding the rules by which differing regions interact, and how they ‘read’ outputs from other regions sending them inputs (‘source transformation rules’). Intracranial recordings of neuronal action potentials have provided great insights into neuronal coding in certain brain regions. In the hippocampal formation, such recordings have disclosed, for example, place, head direction, and grid cells responsive to aspects of 3D space (O’Keefe and Dostrovsky 1971; Taube, Muller, and Ranck 1990; Moser, Kropff, and Moser 2008). At the limit, recording many connected neurons in multiple brain regions provides the ideal approach to inferring neural source transformation rules. However, most recordings to date have used small numbers of electrodes or tetrodes with recordings restricted to perhaps tens of neurons at the most. Recent advances in recording technology – such as Neuropixels – however, potentially allow simultaneous recordings of many more neurons within and between differing brain regions. These advances, might, therefore increase the probability of recording jointly-interacting pairs (or greater) of neurons in differing brain regions, making the detection and isolation of transformation rules between regions more tractable.

Here, using the hypergeometric distribution, and employing anatomically-tractable connection mapping between regions, we derive a method to calculate the probability distribution of ‘recordable’ connections between groups of neurons. This mathematically-derived distribution is validated by Monte Carlo simulations of directed graphs representing the underlying anatomical connectivity structure. We apply this method to simulated graphs with multiple neurons, to analyses based on counts in rat brain regions, and to connection matrices from the Blue Brain model of the mouse neocortex connectome. Overall, we find the probability of simultaneously-recording directly-connected pairs of neurons in vivo in anatomically-connected regions can approach zero with small sample sizes – as is the case with tetrode recordings. In turn, this lack of directly-connected, paired recordings, constrains inferences regarding inter-regional communication. However, alternative approaches, including new recording technologies (such as Neuropixels) and summing neuronal activity over larger scales, offer great promise for identifying inter-regional communication rules.

To help discover these rules, we started by considering a fundamental question: given a random sampling of neurons from multiple anatomically-connected brain regions, what proportion of neurons sampled from a given brain region are likely to receive at least one connection from the neurons sampled from another given brain region? Our analyses underscore the need for such simultaneous recordings, and suggest, via straightforward calculations, where such recordings might be best undertaken to isolate transformation rules: namely, within and between areas for which there is reliable, quantitative data on the anatomical connectivity between regions. Experimental implications of this analysis suggest that multiple, paired, inter-regional recordings in vivo concentrated on circuits and synapses of known connectivity, topography, coding and behavioural correlates will be best suited to understanding changes in neural coding at successive stages of neural information processing.

We focus here on two differing regions for which quantitative anatomical data are available: connected regions of the rat hippocampal formation, and auditory cortex, respectively. For instance, most cells in the rodent hippocampal area CA1 are place cells – they respond to the position in space occupied by the animal on a moment-to-moment basis. CA1 sends the vast proportion of its efferents to the adjacent subiculum, the major output structure of the hippocampus (Witter and Groenewegen 1990; O’Mara et al. 2001), according to a reliable and ordered set of anatomical rules (Amaral, Dolorfo, and Alvarez-Royo 1991; Cembrowski et al. 2018). Yet, subicular neurons display very heterogenous responses (Brotons-Mas et al. 2017) – from no spatial signal whatsoever, to place, grid, boundary-vector and other possible cell classes (of course, the subiculum receives inputs from many other brain regions that might affect neural coding within subiculum; O’Mara 2005). One approach to understanding the rules the subiculum applies to the inputs it receives from area CA1 is from recording in CA1 and subiculum simultaneously, to try and infer from experimental data what these transformation rules might be. Our calculations suggest that there is a high probability of recording multiple directly interacting neurons in anatomically densely-connected regions with high site count neural recording technologies, such as Neuropixels. CA1 is a particularly prominent example of this, given that CA1 has a major output structure to which it sends vast connections; the subiculum. However, this is highly conditional on placing the recording probes in anatomically-appropriate regions of CA1 and subiculum due to the columnar nature of subicular projections from CA1.

Our analyses, based on the Blue Brain Project, suggest that in certain areas of the mouse neocortex, such as the auditory cortex, conjoint recordings of directly-connected or directly-interacting neurons are feasible with new recording technology. By our calculations, when recording 79 neurons uniformly in AUDp (primary auditory area) and AUDpo (posterior auditory area), on average, 30% of neurons in AUDp would receive a direct connection from at least one neuron in AUDpo. Additionally, if connections through multiple synapses are considered (indirect and direct connections), then even for regions with lower connectivity, such as MOp to SSp-ll, the number of connected neurons grows exponentially with the sample rate. This is in line with findings from techniques such as optogenetics, where acting on just a small subset of the full population can have a profound impact.

Combining our work with improvements in connectomics and recording techniques may assist in techniques for analysing neural source transformation rules. For instance, Elsayed and Cunningham (2017) asked if structure in neuronal population recordings is simply to be expected: presenting a statistical method to test if population level results are an expected byproduct of the primary features of temporal and signal correlations and neural tuning. In a similar vein, if our formula indicates that it is very unlikely to record direct connections (or connections with a small number of synapses) between two regions, but high correlation between spiking cells is still found, then it would stand to reason that it has either been observed by chance, or some other region is regulating the activity, or there is a wider oscillatory pattern in the brain driving the activity - but the interacting regions are not directly driving each others activity.

The considerations here allow us to approach the dynamics of, for example, Hebbian theories of learning or synaptic plasticity expressed in neuronal circuits. If, for example, learning is a change in synaptic weights, then, theoretically, at least, we should be able to detect changes in firing rates between interacting neurons embedded in neuronal circuits, even if the sum of the synaptic weights present in the circuit overall do not change. Alternatively, the more elaborate Hebb ‘reverberatory circuit’ view suggests memory should arise as a combination of synaptic weight changes, as well as specific circuit changes or sculpting underly learning and memory. On this latter view, the pattern of activity in the circuit is important, and is sculpted by synaptic weight changes. Conjoint recordings can potentially, therefore, arbitrate on models or theories of synaptic plasticity as a memory mechanism, especially if coupled with other technologies (such as optogenetics).

Overall, we find the probability of simultaneously-recording directly-connected pairs of neurons in vivo in anatomically-connected regions approaches zero unless the regions are well connected and very high site count recording techniques are used. This suggests we may be able to infer source transformation rules from multiple, paired, inter-regional recordings in vivo, in combination with techniques summing activity across many neurons. Additionally, the physical properties of the neural recording device (e.g. tetrodes sample in spheres; probes in capsules; calcium imaging in thin layers) and the angle of approach (i.e. parallel to a columnar structure of cutting across columns) will change the odds of recording connections, emphasising the importance of anatomical sophistication to decide on placements. We envisage that future work would develop statistical models that account for space by considering the neurons in a 3D volume, and then, account for time and dynamics (e.g. firing rates and synaptic strength). The findings presented here will provide a foundation for considering the probability of simultaneously recording connected neurons in multiple, anatomically connected brain regions, and considering pathways to understanding neuronal source transformation rules. Furthermore, they highlight the need to develop analytic techniques to isolate causal structure in such recordings, in order to test theories of synaptic plasticity and memory. Finally, the analyses here underscore the need for further refinement of quantitative digital neuroanatomical techniques to further constrain functional theories of neural coding.

## 4 Methods

### 4.1 Directed graph representation

A directed graph is represented by a sparse matrix indicating the presence of edges in the graph. As such, directed graphs were represented as lists of target nodes, similar to how a sparse matrix can be efficiently stored as a list of lists. For instance, the representation [2, 3], [ ], [0], [1, 2] describes a graph with four vertices 0, 1, 2, 3 and five edges 0 *→* 2, 0 *→* 3, 2 *→* 0, 3 *→* 1, 3 *→* 2. A connection matrix representation of a directed graph can be converted into this format by taking non-zero entries as (*r, c*) and inserting the value *c* into the list *r*.

### 4.2 Finding paths in directed graphs

Given a list of source and target nodes in a directed graph, the number of target nodes reachable from the source nodes was determined by running an iterative deepening depth-limited depth-first search on the transpose graph (in which the direction of arrows are reversed) starting from each target node, to see if any source node was reachable from the target node. This is equivalent to running a search operation on the original graph, and starting from source nodes to see which target nodes are reachable, but speeds up computation by finding predecessors of target nodes as opposed to the successors of source nodes. The result is a list of target nodes that are reachable from one or more source nodes.

### 4.3 Directed graph generation

Directed graphs were randomly generated in Python, while adhering to configured forms of connectivity. To generate a directed graph representing connections between two brain regions, the number of neurons in each region is provided, as well as distributions of forward, recurrent and local connections in each region. Connections were then randomly generated to adhere to these provided distributions. Connections were sampled with replacement, representing the multiple synapses that could be formed between the same pairs of neurons.

### 4.4 Monte Carlo simulations

For directed graphs, random sets of source and target nodes were sampled multiple times and the target nodes reachable from the source nodes were determined. These simulations were performed to evaluate the statistical calculations by comparing the values obtained to those obtained from Monte Carlo simulations.

Additional calculations were performed to estimate the convergence rate of the estimated distribution from these simulations to the true underlying distribution. This was evaluated by checking the smoothness and variance of the simulation result with varying numbers of iterations. This was performed on network graphs, as well as on a simulated problem, whereby there are two bags of balls being drawn from. The first bag contains red and black balls and the number of red balls drawn from the first bag determines the distribution of red and black balls in the second bag. The final result is the number of red balls drawn from the second bag. The statistics of this problem mimic the way the problem is to be solved, but without complications of directed graphs. In all cases, at least 50,000 samples are needed for a good level of convergence in Monte Carlo simulations.

### 4.5 Graph visualisation

Small directed graph visualisations were performed using Networkx (Hagberg, Schult, and Swart 2008). Nodes in the graph are positioned using the Fruchterman-Reingold force-directed algorithm, whereby the edges in the graph are considered as springs holding the nodes close, and the nodes apply a repelling force. The system is then simulated until equilibrium is reached, and the graph is plotted at this point. As such, well connected groups of neurons would be close to each other in the visualization.

Large directed graph visualisations were performed by considering the graph as a connection matrix. In this connection matrix *C*, row *r* and column *c* having the value 1 indicates the existence of a directed connection between node *r* and *c*. The connection matrix is considered in 150*×*150 sized windows, and the percentage of ones in these squares is calculated. The result is then visualisation using a colourmap, from the minimum of 0% representing no connections between the 22,500 pairs of neurons, to the maximum representing the most highly connected block of neurons in the network.

### 4.6 Blue Brain Project model instances

Connection matrices were downloaded from the Blue brain repository (Reimann et al. 2019), version 1.15. Instance 4 ipsilateral and local connections in the right hemisphere of the brain were downloaded for MOp, SSp-ll, VISp, VISl, AUDp, AUDpo, ILA, PL. The data were processed into smaller sparse matrices containing connections, and are stored on the Open Science Framework at https://osf.io/u396f/. These can be loaded using Scipy to handle the npz format files.

### 4.7 Statistics calculation from connection matrices

Given a connection matrix *C* representing the connections between two brain regions, this method calculates the distribution of forward, recurrent and local connections. Firstly, it is assumed that brain region *A* has *N* neurons, and *B* has *M* neurons, the matrix *C* is size (*N* + *M*) *×* (*N* + *M*). Then, the first top left *N × N* part of *C* represents the local connections in *A*, the top right *N × M* part represents the forward connections from *A* to *B*, the bottom left *M × N* part represents the recurrent connections from *B* to *A*, and the bottom right *M × M* part represents the local connections in *B*. The distribution of connections is then calculated by summing the number of ones appearing along rows, and then taking the average of these (excluding 0 for non-local connections), and the number of non-zero rows in each of these four matrices.

### 4.8 Hypergeometric distribution

The hypergeometric distribution describes the probability of obtaining *k* successes in *n* draws without replacement from a population of size *N* with *K* objects that would be considered a success. Let *X* represent the random variable that takes the value of the number of successes drawn from the population, then

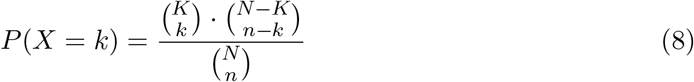

If *N* is much larger than *n*, the binomial distribution can be used instead as a faster approximation of the hypergeometric distribution. In this case, there would be *n* draws from a population of *N* with replacement with *p* = *K/N* as the probability of drawing a success. Then

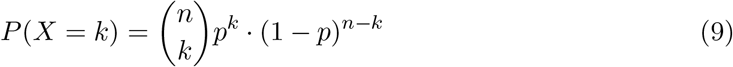

### 4.9 Sum of identically distributed independent random variables

We assume that the random variables *R*_1_, *R*_2_, … , *R*_*k*_ representing the number of synapses that each sampled neuron in *A* that sends connections to *B* are independent and identically distributed with mean *µ* and variance *σ*^2^. We aim is to compute the distribution of the random variable *S*_*k*_ = *R*_1_ + *R*_2_ + … + *R*_*k*_, which represents the number of outgoing synaptic connections from *A*. By the central limit theorem *S*_*k*_ = *R*_1_ + *R*_2_ + … + *R*_*k*_ *≈ N* (*kµ, kσ*^2^), for high values of *k* (usually *k >* 10 is enough for a good approximation, but *k >* 30 is recommended). For *k* below these values, a convolution operation is applied to the distributions of the random variables Grinstead and Snell 2012. If *m*_1_(*x*) and *m*_2_(*x*) are the distribution functions of *X* and *Y*, then the convolution of *m*_1_(*x*) and *m*_2_(*x*) is the distribution function *m*_3_ = *m*_1_ *∗ m*_2_ given by

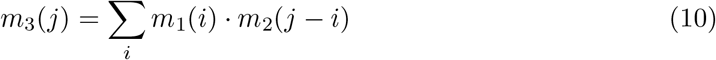

### 4.10 Subsampling distributions

To calculate the probability distribution of the number of sampled neurons that receive connections from another sampled set of neurons can involve very large numbers and distributions. As such, these distributions are often subsampled over a linearly spaced subset to avoid costly calculations. Between the sample points, linear interpolation is performed to estimate the values. However, to improve the accuracy of this operation, the second derivative between the sample points is estimated. This value is compared to that obtained at the next point, and if the difference is larger then the sum of the two values, this indicates that there was a large change in the function between these two sample points. If this is detected, more sample points are evaluated in between these two points to improve the interpolation accuracy.

### 4.11 Python packages used

Many open-source Python packages were vital to this work, and are listed here:

- Mpmath (Johansson and others 2018) - large floating point operations at arbitrary precision and handle cases of non-integer binomial co-efficient calculations, by using the gamma function.
- NumPy (Harris et al. 2020) - random number generation, storage of large arrays, vectorised operations, and gradient estimation.
- Networkx (Hagberg, Schult, and Swart 2008) - infer numerical properties of graphs, and for graph visualisation.
- Scipy (Virtanen et al. 2020) - sparse matrix operations and linear interpolation.
- Matplotlib (Hunter 2007) - plotting colourmaps and graphs.
- Seaborn (Waskom and team 2020) - further plotting operations.
- Pandas (McKinney 2010) - storing and retrieving the results of Monte Carlo simulations.

### Software availability

Python 3.8 code is available, along with instructions to reproduce this work in full and a user interface to run further experiments, from Github at https://github.com/seankmartin/SKMNeuralConnections.

## Acknowledgements

This work was supported by a Joint Senior Investigator Award made by The Wellcome Trust to JPA (103722/Z14/Z) and SMOM.

## Author contributions

SKM: Developing analysis algorithms, Python script writing and validation, analysis and interpretation of data, drafting or revising the article JPA, SMOM: Conception and design, analysis and interpretation of data, drafting and revising the article

## Supplemental material

### S1 Calculations in the hippocampal formation case study

The calculations involved in determining the parameters of the statistical model in the hippocampal formation case study are here. See Table 1 for the numbers that are used in the calculations below. For CA3 to CA1, we calculate, for a tetrode in each region:

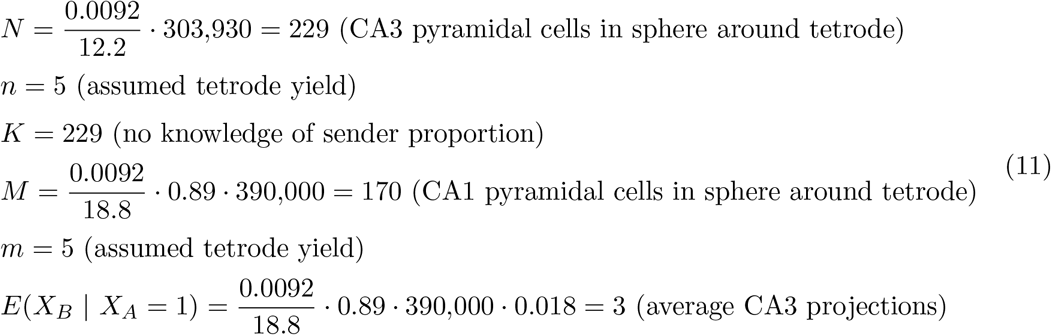

While for a neuropixels probe in each region:

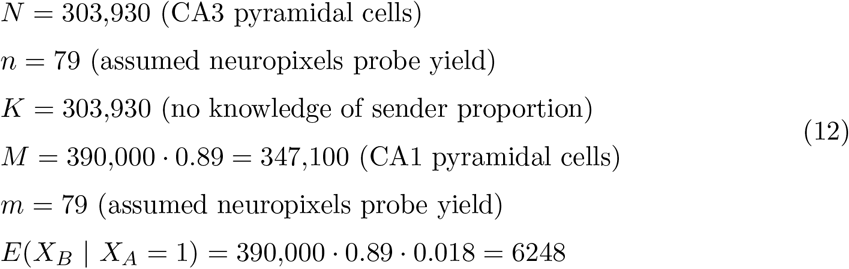

For proximal CA1 to distal subiculum, we calculate, for the high projection rate:

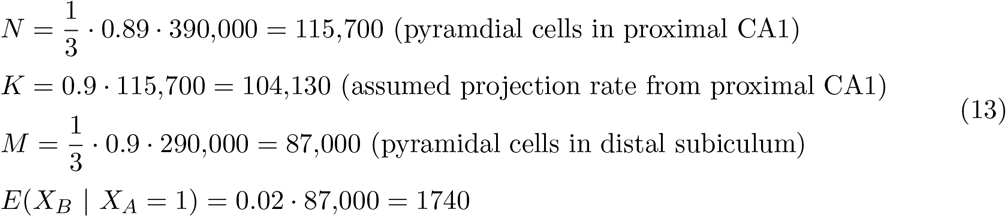

While for the low projection rate:

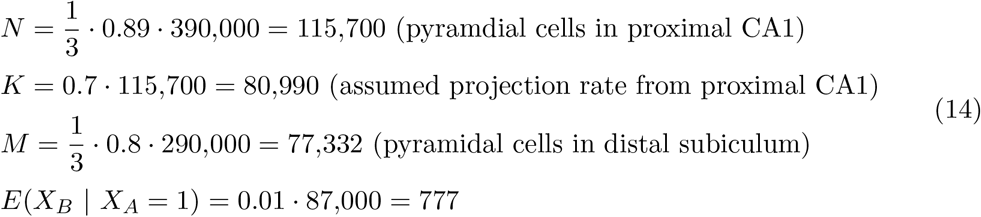

### S2 Accuracy of mean estimation versus full distribution estimation

Using the mean of *P* (*X*_*B*_ = *l* | *X*_*A*_ = *k*) is not as accurate as using the full distribution, but may be required in some cases (e.g. large graphs and indirect connections) for computational performance reasons. Furthermore, in general, the statistical estimation is not as accurate at larger maximum geodesic distances, i.e. indirect connections (see Figure S1).

### S3 Explaining the distributions involved in the statistical estimation

For the graph in Figure 1 panel F, we present the distributions used to obtain the final result of *P* (*X* = *x*). This visualisation may help to understand how the final probability distribution result is obtained (see Figure S2).

### S4 Interpolation accuracy improvement

Since the numbers involved in calculating *P* (*X* = *x*) can be very large, the distributions are sometimes taken at sample points, and linearly interpolated in between these points to avoid calculating the full distribution. This allows the method to be run on larger graphs, without having to use the more inaccurate mean estimation method. To help ensure that important information is not missed, the second derivative is taken in between the points, and a large change in this value indicates missing information. (see Figure S3).

**Figure S1:**
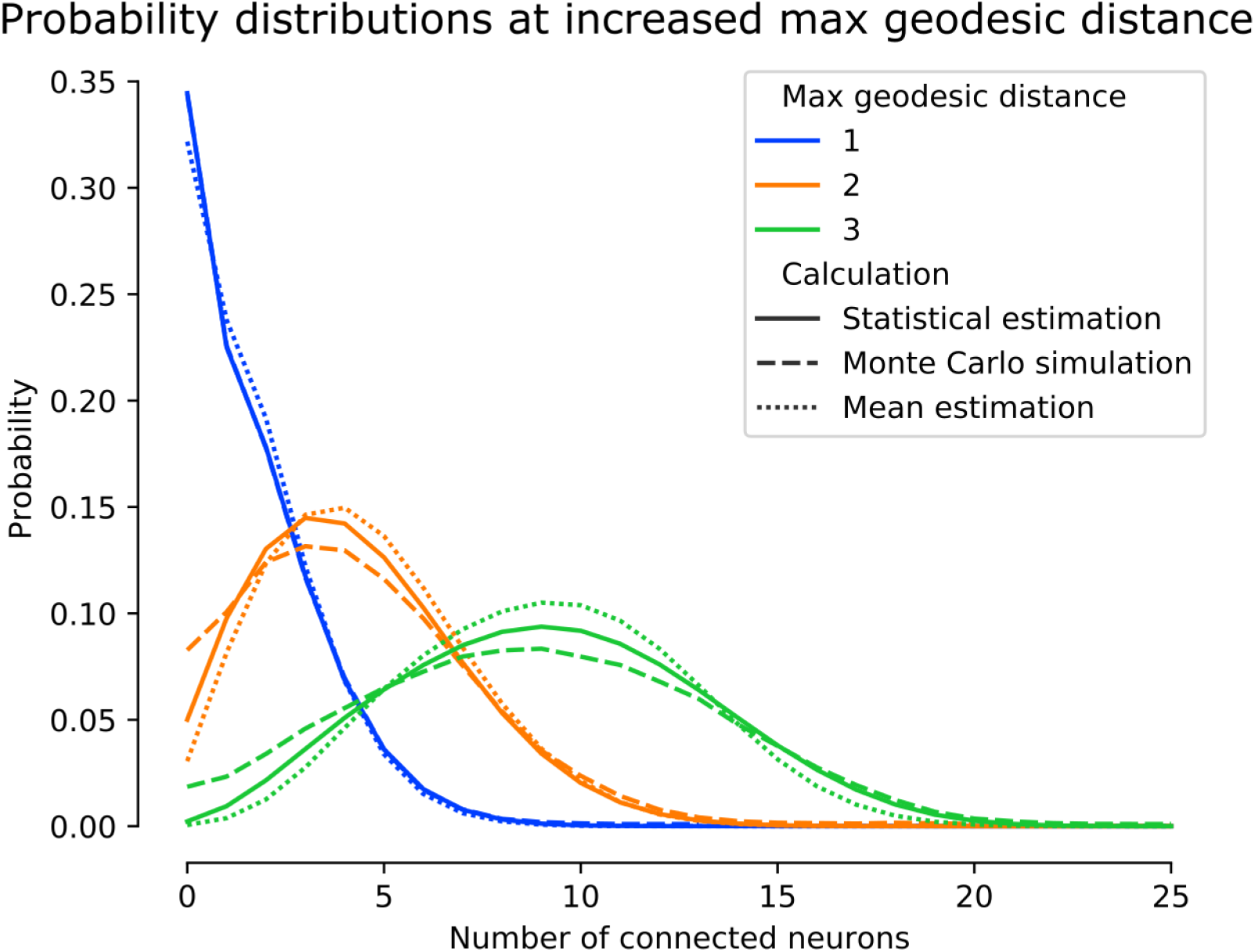
The relationship between accuracy and maximum geodesic distance. As the maximum geodesic distance is increased the accuracy in estimating the probability mass function reduces. However, the expectation of the distribution is a better match. These expectations are; max distance 1: 1.59, 1.59, 1.59, max distance 2: 4.29, 4.33, 4.55, max distance 3: 8.86, 9.27, 9.22, in the order; Monte Carlo simulation, statistical estimation; mean estimation.

**Figure S2:**
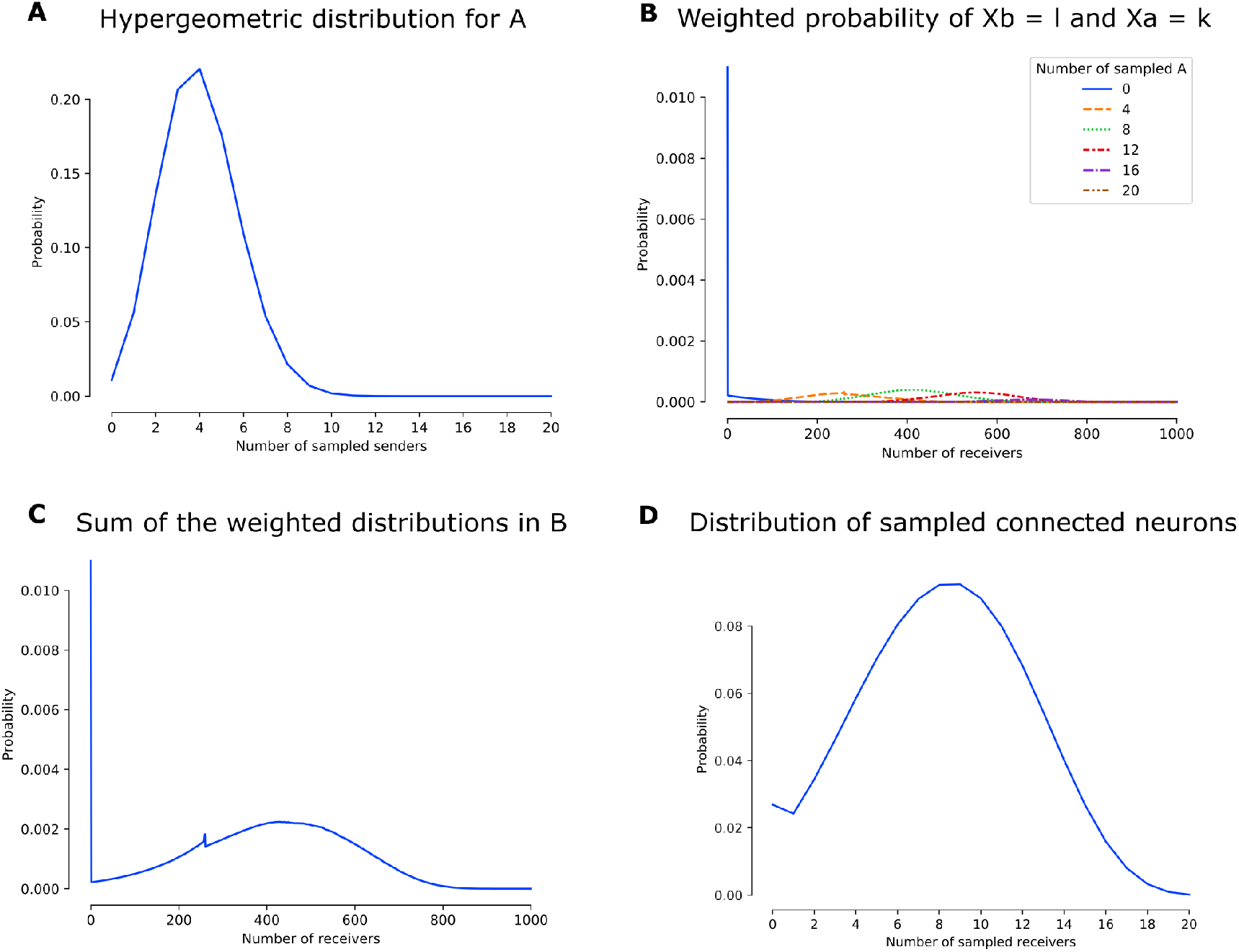
The distributions involved in calculating *P* (*X* = *x*) for Figure 1, F with 20 samples in each region. (A) Plotting *k* against *P* (*X*_*A*_ = *k*), which is hypergeometrically distributed. (B) Plotting *l* against *P* (*X*_*A*_ = *k*) *· P* (*X*_*B*_ = *l* | *X*_*A*_ = *k*) for different values of *k*. (C) Summing the functions in *B* gives the marginal distribution of *P* (*X*_*B*_ = *l*). Note that the blip on the graph around 260 receivers is due to a spike in *P* (*X*_*A*_ = 2) *· P* (*X*_*B*_ = *l* | *X*_*A*_ = 2) around *l* = 260, followed by a sharp decrease. (D) The final result of *x* against *P* (*X* = *x*) by using the distribution in C to calculate *P* (*X* = *x* | *X*_*B*_ = *l*) *· P* (*X*_*B*_ = *l*) and then summing over *l* to obtain the final distribution.

**Figure S3:**
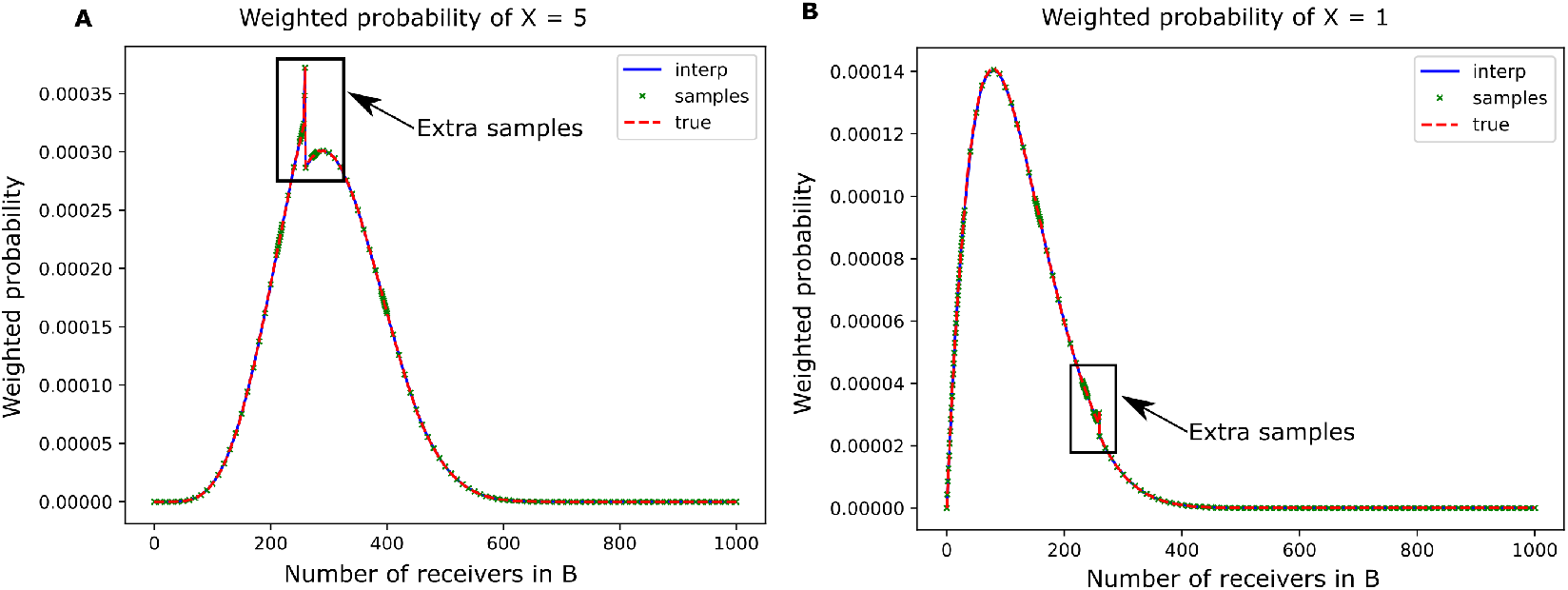
The second derivative is used to refine samples and find large change points. (A) Plotting *x* against *P* (*X* = 5 |*X*_*B*_ = *x*) *· P* (*X*_*B*_ = *x*). Extra samples are required for an accurate subsampling that would not be picked up without running a detection step for changes. (B) As in (A), but plotting *x* against *P* (*X* = 1 |*X*_*B*_ = *x*) *· P* (*X*_*B*_ = *x*).

